# The Tension-sensitive Ion Transport Activity of MSL8 is Critical for its Function in Pollen Hydration and Germination

**DOI:** 10.1101/084988

**Authors:** Eric S. Hamilton, Elizabeth S. Haswell

## Abstract

All cells respond to osmotic challenges, including those imposed during normal growth and development. Mechanosensitive (MS) ion channels provide a conserved mechanism for regulating osmotic forces by conducting ions in response to increased membrane tension. We previously demonstrated that the MS ion channel MscS-Like 8 (MSL8) is required for pollen to survive multiple osmotic challenges that occur during the normal process of fertilization, and that it can inhibit pollen germination. However, it remained unclear whether these physiological functions required ion flux through a mechanically gated channel provided by MSL8. We introduced two point mutations into the predicted pore-lining domain of MSL8 that disrupted normal channel function in different ways. The Ile711Ser mutation increased the tension threshold of the MSL8 channel while leaving conductance unchanged, and the Phe720Leu mutation severely disrupted the MSL8 channel. Both of these mutations impaired the ability of MSL8 to preserve pollen viability during hydration and to maintain the integrity of the pollen tube when expressed at endogenous levels. When overexpressed in a *msl8-4* null background, MSL8^I711S^ could partially rescue loss-of-function phenotypes, while MSL8^F720L^ could not. When overexpressed in the wild type L*er* background, MSL8^I711S^ suppressed pollen germination, similar to wild type MSL8. In contrast, MSL8^F720L^ failed to suppress pollen germination and increased pollen bursting, thereby phenocopying the *msl8-4* mutant. Thus, an intact MSL8 channel is required to for normal pollen function during hydration and germination. These data establish MSL8 as the first plant MS channel to fulfill previously established criteria for assignment as a mechanotransducer.

## Introduction

In order to thrive, all cells must respond to osmotic challenges. Drought, salt stress, and freezing all present environmentally imposed stresses that require osmoprotective strategies for cell survival (Burg and Ferraris, 2008; Zhang, 1999). Organisms can also experience osmotic challenges inherent to their growth and development, such as desiccation during endospore formation in response to stress (Tovar-Rojo et al., 2003), the transition from seawater to saltwater experienced by migrating salmon (Jeffries et al., 2011), or drying and rehydration of plant seeds (Hoekstra et al., 2001).

One conserved molecular mechanism for responding to osmotic challenges and other mechanical stimuli is the use of mechanosensitive (MS) ion channels (Booth and Blount, 2012; Hamilton et al., 2015b; Ranade et al., 2015). These proteins form pores in the cell membrane that open in response to mechanical stimulation, allowing ions to flow across the membrane down their electrochemical gradient. A role for MS ion channels in osmotic regulation has been well described in *Escherichia coli*, where the Mechanosensitive channel of Large conductance (MscL) and Mechanosensitive channel of Small conductance (MscS) are required for the cell to survive extreme hypoosmotic downshock, such as the transfer from 500 mM NaCl to distilled water (Levina et al., 1999). It is proposed that when cells are exposed to hypoosmotic shock, the resultant cell swelling and increase in membrane tension increase the open probability of both MscS and MscL. When open, these channels allow osmolytes to flow out of the cell, reducing internal osmotic pressure and protecting the bacterium from lysis (Booth and Blount, 2012; Kung et al., 2010). Thus, bacterial MS ion channels are often referred to as osmotic pressure release valves.

*Ec*MscS is the founding member of a large family of conserved proteins found within prokaryotes, archaea, and many eukaryotes, including plants (Haswell, 2007; Kloda and Martinac, 2002; Koprowski and Kubalski, 2003; Levina et al., 1999; Malcolm and Maurer, 2012; Pivetti et al., 2003; Porter et al., 2009; Prole and Taylor, 2013). Crystal structures of MscS homologs from *E. coli* and *Helicobacter pylori* have shown that these proteins form homoheptamers (Bass et al., 2002; Lai et al., 2013; Steinbacher et al., 2007; Wang et al., 2008), but no structural information yet exists for eukaryotic MscS family members. Ten MscS homologs are encoded in the *Arabidopsis thaliana* genome, named MscS-Like (MSL) 1-10 (Haswell, 2007). MSL proteins localize to diverse cell compartments, including the mitochondria (Lee et al., 2016), plastids (Haswell and Meyerowitz, 2006), and the plasma membrane (Hamilton et al., 2015a; Haswell et al., 2008; Veley et al., 2014).

We recently showed that plasma membrane-localized MSL8 is required to protect pollen from osmotic challenges during several steps in fertilization (Hamilton et al., 2015a). Pollen contains the male gametes of flowering plants and is responsible for the delivery of two sperm cells to the female gametophyte in order to produce the next generation (Hafidh et al., 2016). During key steps of desiccation, rehydration, germination, and tube growth, pollen must regulate its osmotic potential to maintain the integrity of the cell and achieve its reproductive function (Beauzamy et al., 2014; Chen et al., 2015; Feijo et al., 1995; Firon et al., 2012; Sanati Nezhad et al., 2013).

Upon maturation, the pollen of most species of flowering plants desiccates (Franchi et al., 2011), a strategy thought to maintain viability during exposure to the external environment (Firon et al., 2012). To protect against the osmotic stress of this water loss, pollen accumulates sugars and other compatible osmolytes that serve to stabilize cellular membranes and other biomolecules in the dry state (Hoekstra et al., 2001; Pacini et al., 2006). When a desiccated pollen grain reaches the stigma, an organ of finger-like projections atop the pistil, it is rehydrated by stigma cell exudate in a regulated process of pollen reception (Dresselhaus and Franklin-Tong, 2013; Edlund, 2004; Samuel et al., 2009). The stigma exudate of Lily, which has a wet stigma coated in exudate, is more than 90% water, with carbohydrates making up the majority of the solutes (Labarca et al., 1970). The composition of the exudate produced in species with dry stigmas, such as Arabidopsis, is unknown (Edlund, 2004), but must be hypoosmotic with respect to the pollen cytoplasm in order to produce net hydration. The rehydration of desiccated pollen can damage dry cellular membranes, which are more rigid and less able to accommodate hypoosmotic swelling (Hoekstra et al., 2001; 1997), and this membrane damage can compromise pollen viability if hydration is not properly regulated (van Bilsen et al., 1994).

Once a pollen grain is rehydrated, its metabolism is reactivated and it germinates a long extension that grows by turgor-driven cell expansion at the tip without further cell divisions, called a pollen tube (Beauzamy et al., 2014; Firon et al., 2012). Germination requires that the pollen grain establish and maintain turgor pressure at a level high enough to break through the tough pollen cell wall (Feijo et al., 1995). The rapid growth of the pollen tube is also driven by turgor pressure, which continuously expands new cell wall material delivered to the growing tip (Hill et al., 2012; Kroeger et al., 2011; Zerzour et al., 2009). Artificially increasing turgor pressure can lead to lysis, especially at the tip of the pollen tube (Benkert et al., 1997), and genetic lesions that increase turgor impair the ability of the pollen tube to reach the ovule and fertilize the egg cell (Chen et al., 2015). Conversely, growth conditions that reduce turgor halt pollen tube growth (Zerzour et al., 2009). In addition to turgor, pollen tubes must regulate the delivery of new cell wall material and cell wall composition in order to sustain growth while maintaining the integrity of the cell (Hill et al., 2012; Kroeger et al., 2011; Zerzour et al., 2009). These observations have led to the proposal that pollen grains utilize a mechanosensor to respond to changes in osmotic potential (Hill et al., 2012), and a role for MS ion channels in regulating ion fluxes at the dynamic pollen tube tip has been proposed for decades (Feijo et al., 1995).

MSL8 provides one mechanism for regulating these osmotic forces in pollen. Both loss of function mutations in the *MSL8* gene and *MSL8* overexpression result in male fertility defects, indicating that the proper expression level of *MSL8* is essential for full reproductive success (Hamilton et al., 2015a). Two independent T-DNA insertion alleles, the partial loss-of-function *msl8-1* in the Col*-0* background and the null *msl8-4* in the L*er* background, result in reduced viability when pollen is hydrated in distilled water. This hydration viability defect can be rescued by supplementing hydration media with a 20% solution of polyethylene glycol (PEG); increasing the osmotic potential of the media reduces the hypoosmotic shock pollen experiences during hydration. Expression of *MSL8-GFP* from genomic sequences fully complements this reduced viability during *in vitro* hydration in both allele backgrounds. MSL8 also negatively regulates pollen germination while maintaining the integrity of germinating pollen tubes. Mutant *msl8* pollen germinates at a greater rate than wild type pollen, but bursts more frequently. Overexpressing *MSL8* in the L*er* background strongly suppresses pollen germination, which leads to reduced fertility.

Thus, MSL8 is required to protect pollen against excessive osmotic pressure, either during rehydration or during the turgor-driven germination and growth of a pollen tube. Because MSL8 forms a MS ion channel, we hypothesized that the flow of ions through the MSL8 pore in response to increases in membrane tension is critical for preventing lysis during hydration and germination, akin to the function of MscS in *E. coli*. However, it remained possible that the role of MSL8 in pollen survival is indirect. Indeed, close homolog MSL10 triggers programmed cell death signaling independent of its channel activity (Veley et al., 2014).

Here we report the results of experiments aimed to determine if MSL8 requires an intact channel to achieve its function in pollen. To do so, we mutated the predicted pore of the MSL8 channel, identified lesions that affect channel behavior, and analyzed the ability of these mutants to function in pollen. Our results support a model wherein MSL8 regulates osmotic forces in pollen directly by transporting ions.

## Results

### Mutations in the predicted pore-lining domain of MSL8 alter its channel properties

In order to alter the channel properties of MSL8, we identified candidate amino acids for mutagenesis based alignment with *Ec*MscS. The predicted topology of a MSL8 monomer shown in Fig. 1A indicates a protein with 6 transmembrane (TM) helices and three soluble domains. The most C-terminal TM helix, TM6, contains sequence with modest homology to the pore-lining TM3 of *Ec*MscS (marked with a thick line in Fig. 1A and sequence shown in Fig. 1B).

**Figure 1.**
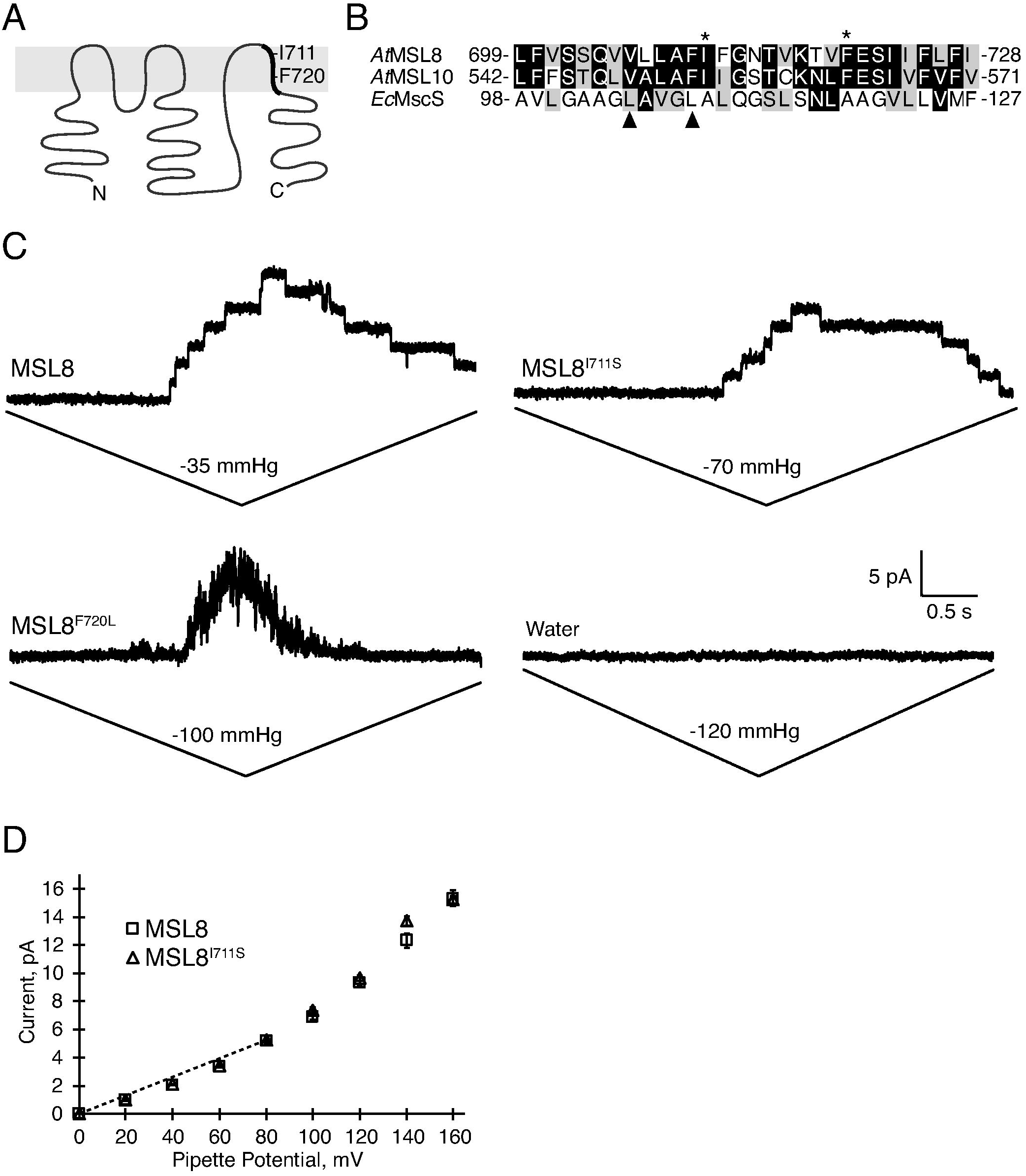
Mutations in the predicted pore-lining domain alter channel properties of MSL8. (**A**) Predicted topology of MSL8. N and C mark the amin– and carboxy-terminal ends respectively. Thick line marks the predicted pore-lining domain. Residues mutated in this study are indicated. (**B**) Multiple alignment of the predicted pore-lining transmembrane regions of Arabidopsis MSL8, Arabidopsis MSL10 and the known pore-lining domain of *E. coli MscS*. Identical residues conserved in at least half of the sequences are shaded darkly; similar residues conserved at this level are shaded in gray. Residues proposed to form the channel seal in *Ec*MscS are marked with arrowheads. MSL8 residues mutated in this study are marked with an asterisk. (**C**) Representative traces from excised inside-out patches of plasma membrane from *Xenopus laevis* oocytes following injection with the indicated cRNA or water clamped at -40 mV membrane potential. (**D**) The current-voltage relationship of MSL8 (squares, *N* = 4 oocytes) and MSL8^I711S^ (triangles, *N* = 9 oocytes) in symmetric 60 mM MgCl_2_. Dashed line indicates the slope from which the single-channel conductance was measured (see Table 1). Error bars are mean ± SE.

In the *Ec*MscS channel, Leu105 and Leu109 from each of the seven monomers are proposed to form a hydrophobic seal in the closed channel that prevents the flow of ions across the pore (Anishkin and Sukharev, 2004; Bass et al., 2002; Steinbacher et al., 2007) (arrowheads, Fig. 1B). Mutations in nearby residues can result in decreased tension sensitivity, increased tension sensitivity, or reduced conductance (Belyy et al., 2010; Edwards et al., 2005; Rasmussen et al., 2015). To create MSL8 variants with altered channel properties, we replaced hydrophobic residues surrounding the presumptive hydrophobic seal with polar residues or smaller non-polar residues. Two mutants, Ile711 to Ser and Phe720 to Leu (asterisks, Fig. 1B) were selected for further study based on their ability to produce MS currents when expressed in *Xenopus laevis* oocytes (see below).

*MSL8*, *MSL8^I711S^* and *MSL8^F720L^* cRNAs were injected into *Xenopus* oocytes for analysis by single channel patch-clamp electrophysiology as previously described (Hamilton et al., 2015a; Maksaev and Haswell, 2015). Patches of plasma membrane were excised from oocytes 2 to 10 days after injection, and current measured over time as membrane tension in the patch was increased by applying suction through the pipette using a pressure-clamp controller. In symmetric 60 mM MgCl_2_ + 5mM HEPES, oocytes expressing wild type *MSL8* cRNA exhibited tension-gated activity as expected (Fig. 1C, top left panel), as did oocytes injected with *MSL8^I711S^* cRNA (Fig. 1C, top right panel). In the same system, *MSL8^F720L^* cRNA produced MS currents without discrete channel openings (Fig. 1C, bottom left panel; see Supplementary Fig. S1 for more examples of activity associated with *MSL8^F720L^* cRNA), and no currents were observed in water-injected oocytes (Fig. 1C, bottom right panel). MS currents were observed in 23 of 26 patches pulled from oocytes injected with *MSL8* cRNA; 27 of 32 patches for *MSL8^I711S^*; 20 out of 24 patches for *MSL8^F720L^*; and in none of 26 patches from water-injected oocytes (Table 1). Under these conditions, the single-channel conductance of MSL8^I711S^ was indistinguishable from that of wild type MSL8 (Fig. 1D, Table 1). Without distinguishable individual gating events, we were unable to assess the single channel conductance of MSL8^F720L^.

**Table 1.** Conductance, tension threshold, and number of observations of MSL8 variants in *Xenopus* oocytes.

|  | Conductance | Tension Threshold |  | Patches with activity / total patches |
| --- | --- | --- | --- | --- |
|  |  | First channel to open, mmHg | Last channel to close, mmHg |  |
| MSL8 | 60.0 | -18.9 ± 13.1 | -1.7 ± 3.5 | 23/26 |
| MSL8<sup>I711S</sup> | 60.7 | -29.5 ± 11.7 | -9.4 ± 9.5 | 27/32 |
| MSL8<sup>F720L</sup> | NA | -61.2 ± 12.3 | -52.7 ± 11.5 | 20/24 |
| Water | NA | NA | NA | 0/26 |

We were unable to calculate a midpoint gating tension, because membrane patches ruptured before current saturation, as previously observed for MSL10 expressed in *Xenopus* oocytes (Maksaev and Haswell, 2012). Instead, we estimated the tension sensitivity of each MSL8 variant, by recording the suction associated with the opening of the first channel and the closing of the last channel in each patch. Patch pipettes of size 3 ± 0.5 MOhm were used in all experiments to reduce variability in patch geometry. The negative pressure required to open the first channel in patches expressing wild type MSL8 was -18.0 ± 13.1 mmHg, while for MSL8^I711S^ it was -29.5 ± 11.7 mmHg (Table 1). While we were unable to identify discrete channel openings for MSL8^F720L^, we estimated its tension threshold by determining the pressure required to produce visible MS currents, on average -61.2 ± 12.3 mmHg (Table 1). In summary, MSL8^I711S^ had the same conductance as wild type MSL8, but was significantly less tension-sensitive (required greater negative pressure to open). MSL8^F720L^ appears to be a severely disrupted channel, albeit one capable of conducting ions through its pore in response to extremely high membrane tension, approximately 3 times higher than that required to open wild type MSL8.

### MSL8 channel mutants do not complement msl8-4 mutant phenotypes when expressed from genomic sequences

To determine whether channel activity was required for MSL8 to protect pollen from osmotic challenges, we tested whether MSL8^I711S^ or MSL8^F720L^ could complement previously characterized *msl8-4* mutant defects. MSL8, MSL8^I711S^ and MSL8^F720L^ were tagged with GFP at the C-terminus and expressed from genomic sequences in the null *msl8-4* mutant background. The wild type version of this transgene, termed *gMSL8-GFP*, was shown to rescue the *in vitro* hydration viability defect of *msl8-4* pollen (Hamilton et al., 2015a). Lines expressing wild type and mutant versions of *gMSL8-GFP* were generated and screened for expression of *MSL8* variants in floral RNA. Two independent homozygous transgenic lines expressing *gMSL8^I711S^-GFP* or *gMSL8^F720L^-GFP* and a single control line expressing wild type *gMSL8-GFP* were selected for further analysis. All lines selected accumulated *MSL8* transcripts at levels equal to or greater than the wild-type L*er* (Fig. 2A). To confirm protein production, stability and subcellular localization, confocal images were taken of GFP signal in mature, desiccated pollen isolated from these lines. All five lines expressing *gMSL8-GFP* or variants in the *msl8-4* background exhibited GFP signal localized to both the plasma membrane and endomembrane compartments and excluded from vacuoles, as previously observed for MSL8-GFP (Fig. 2B; (Hamilton et al., 2015a). Untransformed *msl8-4* mutant pollen only exhibited autofluorescence of the cell wall.

**Figure 2.**
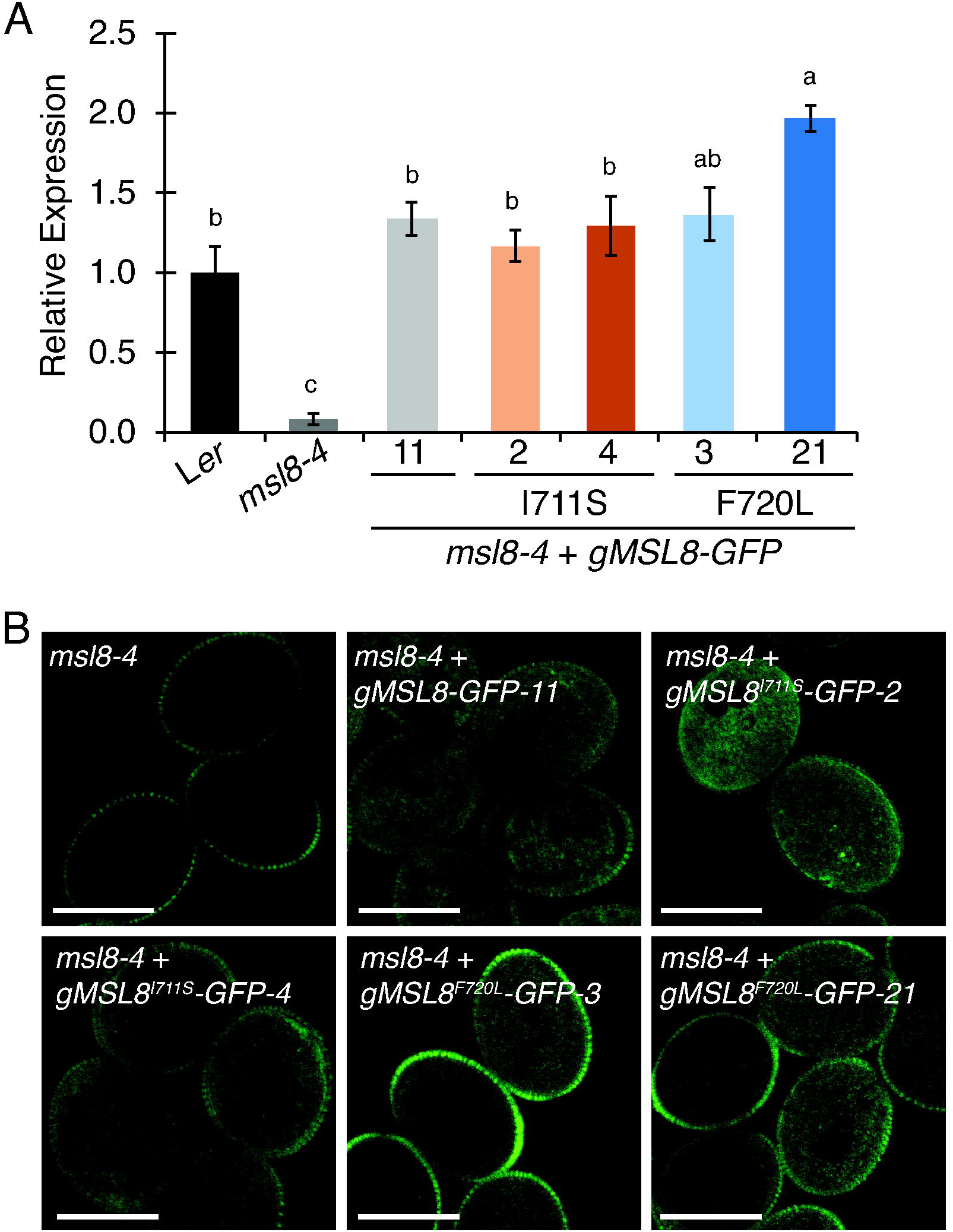
Expression and subcellular localization of MSL8-GFP expressed from endogenous sequences. **A**) Quantitative reverse-transcription polymerase chain reaction amplification of *MSL8* transcripts in L*er*, *msl8-4*, and *gMSL8-GFP* transgenic lines. Levels are presented relative to *ACTIN*. Different letter groups indicate significant (p < 0.05) differences between groups as determined by Tukey’s post-hoc test following one-way ANOVA. Error bars are mean ± SE of three biological and two technical replicates. (**B**) Confocal images of GFP signal in pollen from stage 13-14 flowers from the indicated lines hydrated in water. Scale bar is 20 μm.

To test the ability of MSL8^I711S^ and MSL8^F720L^ to protect *msl8-4* pollen from the hypoosmotic shock of rehydration, we quantified the viability of mature, desiccated pollen after rehydration in distilled water. In a rehydration time course from 30 to 120 minutes, As expected, L*er* pollen viability exceeded 80% at all time points, while *msl8-4* pollen exhibited less than 35% survival on average, and *msl8-4* pollen expressing wild type *gMSL8-GFP* exhibited wild-type survival at all time points (Fig. 3A). However, there were no differences between the viability of *msl8-4* pollen and *msl8-4* pollen expressing the *gMSL8^I711S^-GFP* or *gMSL8^F720L^-GFP* transgenes in water at 30, 60, or 120 minutes after hydration. Thus, neither MSL8^I711S^ nor MSL8^F720L^ could complement this *msl8-4* mutant phenotype.

**Figure 3.**
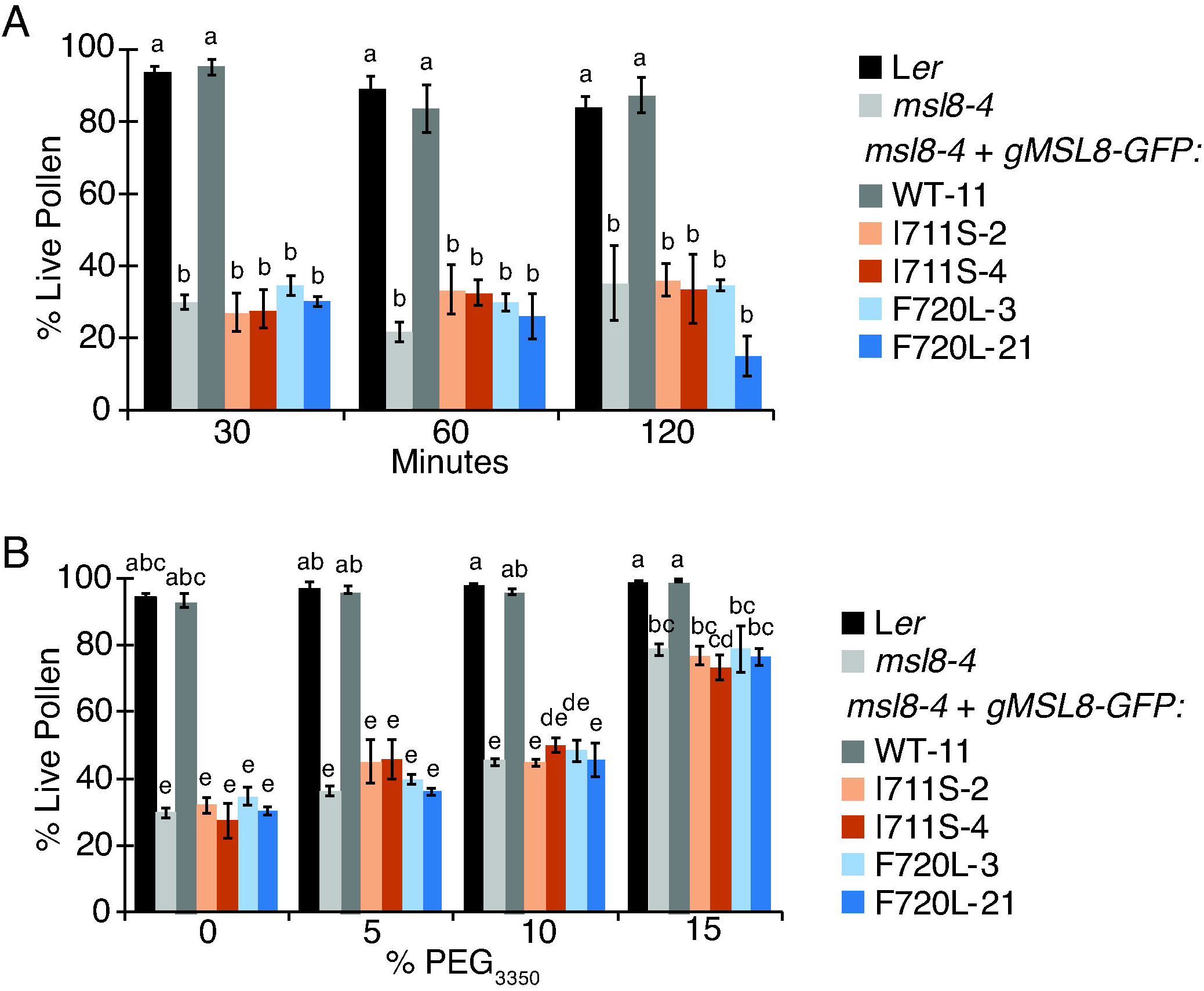
MSL8^I711S^-GFP and MSL8^F720L^-GFP do not complement the *msl8-4* hydration viability defect when expressed from endogenous sequences. (**A**) Hydration viability time course. Mature, desiccated pollen from *msl8-4* + gMSL8-GFP and variant lines hydrated in distilled water for the indicated periods of time. Viability was determined by staining with fluorescein diacetate (which marks live pollen) and propidium iodide (which enters compromised cells). The average of 3-6 experiments with *N* = 58 – 377 pollen grains per experiment is shown. (**B**) Hydration viability in PEG3350 series. The average of 3-9 experiments with *N* = 68-566 pollen grains per experiment is presented. Different letter groups indicate significant (p < 0.05) differences between groups as determined by Scheffe’s post-hoc test following tw–way ANOVA. Error bars are mean ± SE.

To determine if MSL8^I711S^ or MSL8^F720L^ could protect *msl8-4* pollen from intermediate levels of hypoosmotic stress, we quantified pollen viability following a 30-minute incubation in a range of PEG_3350_ concentrations. Supplementing hydration media with 15% PEG_3350_ increased the average viability of *msl8-4* pollen from 30% to 79% (Fig. 3B). PEG supplementation also increased the survival rate of *msl8-4* pollen expressing *gMSL8^I711S^-GFP* and *gMSL8^F720L^-GFP*, to a degree that was indistinguishable from that of the mutant *msl8-4* pollen across all levels of PEG (Fig. 3B). Thus, the *gMSL8^I711S^-GFP* and *gMSL8^F720L^-GFP* transgenes did not alter the sensitivity of *msl8-4* pollen to rehydration, even when osmotic supplementation was used to reduce the severity of the osmotic downshock.

*msl8-4* pollen survives hydration in germination media, as it contains osmolytes (580 mOsmol), but exhibits increased germination rates and pronounced bursting when germinated *in vitro* (Hamilton et al., 2015a). We therefore quantified the germination and bursting rates of *msl8-4* pollen expressing *gMSL8^I711S^-GFP*- and *gMSL8^F720L^-GFP*. After incubation in germination media for 6 hours, most L*er* pollen was ungerminated and intact; 8% was ungerminated and burst; 12% germinated and was intact; and 4% germinated and burst (Fig. 4, Supplementary Table S1). *msl8-4* pollen germinated at a similar rate to L*er*, but 37% of ungerminated mutant pollen grains burst. Most of the *msl8-4* pollen that did germinate went on to burst, leaving only 49% intact pollen (compared to 89% for the wild type). While *msl8-4* pollen expressing *gMSL8-GFP* barely germinated or burst at all (< 0.5%) under these conditions, pollen expressing *gMSL8^I711S^-GFP* or *gMSL8^F720L^-GFP* had germination and bursting rates similar to *msl8-4* pollen. We thus found no evidence that MSL8^I711S^ or MSL8^F720L^ could substantially rescue known *msl8-4* loss-of-function phenotypes (Figs. 3-4), even though these variants were expressed at or above native levels and localized normally (Fig. 2).

**Figure 4.**
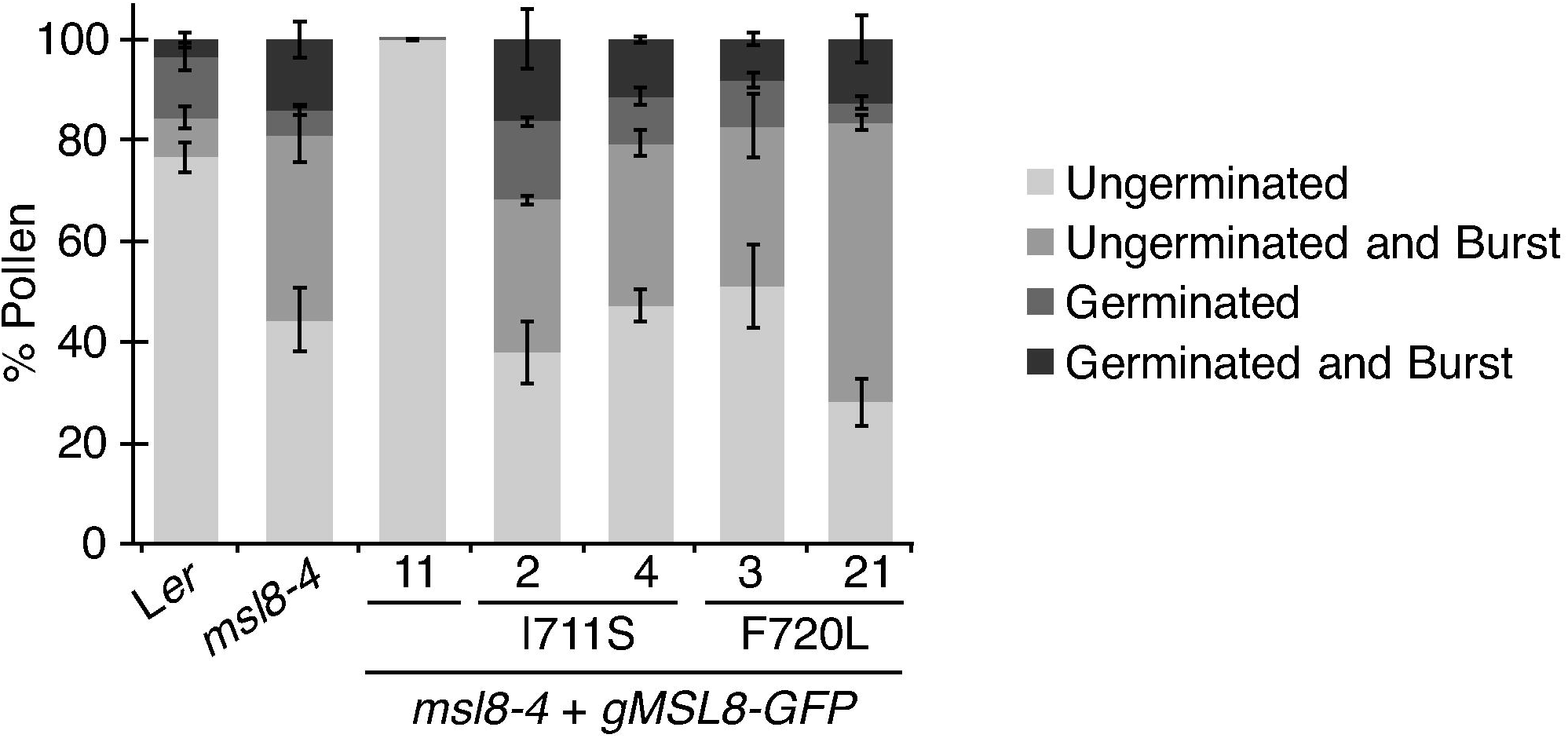
MSL8^I711S^-GFP and MSL8^F720L^-GFP do not suppress *msl8-4* pollen bursting or germination when expressed from endogenous sequences. Mature, desiccated pollen from the indicated lines were incubated in germination media for 6 hours then examined under the microscope. Each pollen grain scored was placed into one of the four indicated categories. Pollen was counted as germinated if it had produced a pollen tube longer than the pollen grain. Pollen was counted as burst if expelled cytoplasm was visible. Averages from 3-6 experiments with *N* = 101-236 pollen per experiment are presented. See Supplementary Table 1 for statistical differences between groups. Error bars are mean ± SE.

*MSL8^I711S^ but not MSL8^F720L^ partially rescues msl8-4 loss-of-function phenotypes when overexpressed.* To determine if we could detect partially functional channels if they were present at high levels, MSL8, MSL8^I711S^ or MSL8^F720L^ were tagged at the C-terminus with YFP and expressed under the control of the strong, pollen-specific promoter *LAT52* in the *msl8-4* background. We identified multiple independent homozygous lines that exhibited a range of transgene expression levels in floral RNA, from 1.9- to 9.6- fold over the levels of endogenous *MSL8* in L*er* (Fig. 5A). YFP signal in *msl8-4* pollen expressing *LAT52pMSL8-YFP* and variants localized to the plasma membrane and endomembrane compartments of mature pollen (Fig. 5B; Supplementary Fig. S2A), and the intensity of YFP signal correlated with transcript level as revealed by RT-PCR (compare Fig. 5A and Supplementary Fig. S2A).

**Figure 5.**
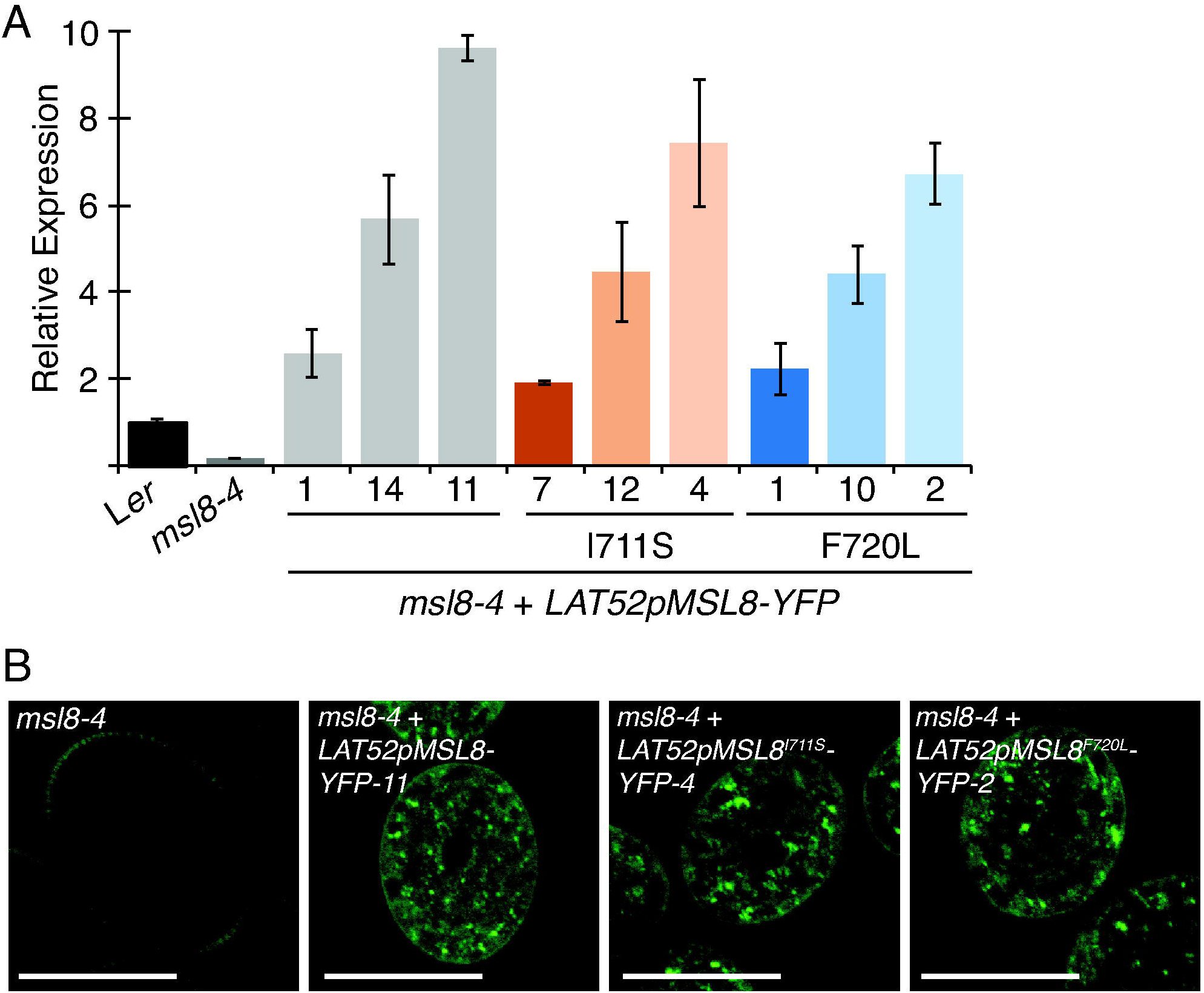
Expression and subcellular localization of MSL8^I711S^-YFP and MSL8^F720L^-YFP expressed from the *LAT52* promoter in the *msl8-4* background. (**A**) Quantitative reverse-transcription polymerase chain reaction amplification of *MSL8* transcripts, presented relative to *ACTIN*, in L*er*, *msl8-4*, and *msl8-4* + *LAT52pMSL8-YFP* transgenic lines. Error bars are mean ± SE. (**B**) Confocal images of YFP signal in *msl8-4* pollen or *msl8-4* pollen expressing *LAT52pMSL8-YFP*, *LAT52pMSL8^I711S^-YFP*, or *LAT52pMSL8^F720L^-YFP*. Scale bars are 20 μm.

Pollen from L*er* and *msl8-4* lines overexpressing MSL8, MSL8^I711S^, and MSL8^F720L^ was incubated in distilled water for 30 or 120 minutes and viability quantified as in Fig. 3. All three *msl8-4* + *LAT52pMSL8-YFP* lines survived as well as wild-type pollen at both time points, while the viability of *msl8-4* pollen expressing *LAT52pMSL8^F720L^-YFP* was indistinguishable from that of *msl8-4* in all three independent lines (Fig. 6). Thus, even at high levels of expression, *MSL8^F720L^* was unable to protect pollen from the hypoosmotic shock of rehydration. However, pollen from three *msl8-4* + *LAT52pMSL8^I711S^-YFP* lines exhibited intermediate levels of pollen viability when hydrated in water. The level of protection was correlated with the level of *MSL8^I711S^* expression in each line. Two independent lines, I711S-7 and I711S-12, exhibited averages of 44% to 63% survival during the time course. At both time points, I711S-7- and I711S-12 pollen viability was statistically significantly different from L*er*, *msl8-4*, and the other transgenic lines, representing an intermediate phenotype. The strongest-expressing line, I711S-4 (Fig. 5A), exhibited wild-type pollen viability at both 30 and 120 minutes. Thus, MSL8^I711S^ is able to partially rescue the *msl8-4* hydration viability defect when moderately overexpressed, while sufficiently high levels of expression fully complement the mutant phenotype.

**Figure 6.**
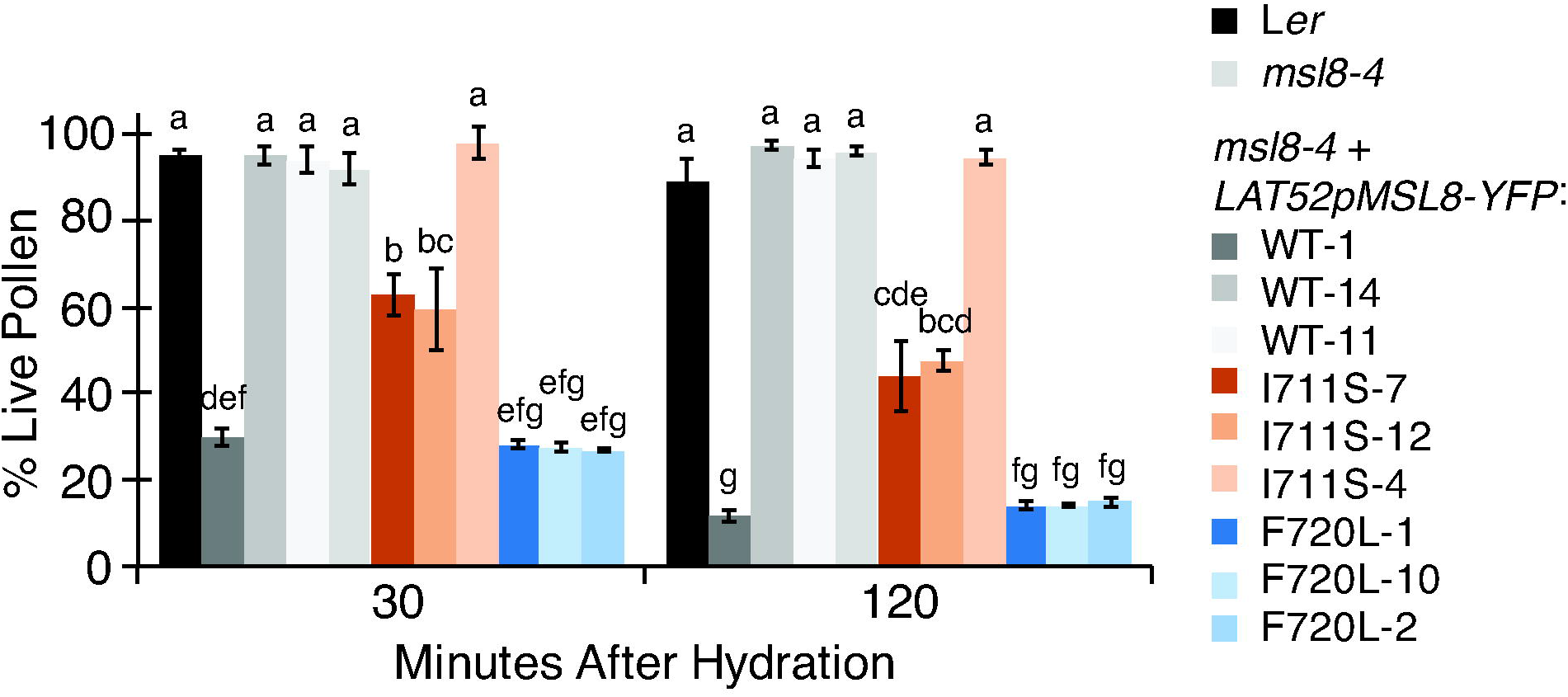
MSL8^I711S^-YFP but not MSL8^F720L^-YFP partially rescues the *msl8-4* hydration viability defect when overexpressed in the *msl8-4* background. Hydration viability time course, performed as described in Fig. 3. The average of 3 to 4 experiments with *N* = 124 to 354 pollen per experiment is presented. Different letter groups indicate significant (p < 0.05) differences between groups as determined by Scheffe’s post-hoc test following two–way ANOVA. Error bars are mean ± SE.

Next, we incubated pollen from *msl8-4* + *LAT52pMSL8-YFP* and variant lines in germination media for 6 hours and quantified germination and bursting rates (Fig. 7A). As expected, expression of *LAT52pMSL8-YFP* suppressed both germination and bursting rates of *msl8-4*, to less than 1%, while expression of *LAT52pMSL8^F720L^-YFP* did not (Fig. 7A, Supplementary Table S2). As with pollen rehydration, overexpression of *MSL8^I711S^* in the *msl8-4* background resulted in partial rescue of the loss-of-function phenotypes. Mutant pollen expressing *LAT52pMSL8^I711S^-YFP* germinated an average of 0% to 17%, and burst on average 4% to 36% of the time (much lower than the approximately 25% germination and 50% bursting of *msl8-4*). Both germination and bursting rates were inversely correlated to the level of *MSL8* accumulation in the *LAT52pMSL8^I711S^-YFP* line (compare Figs. 5A and 7A). The pattern of bursting and germination in the highest expressing line, I711S-4, was statistically indistinguishable from line WT-11 (Supplementary Table S2), indicating that MSL8^I711S^ can fully complement when highly overexpressed.

**Figure 7.**
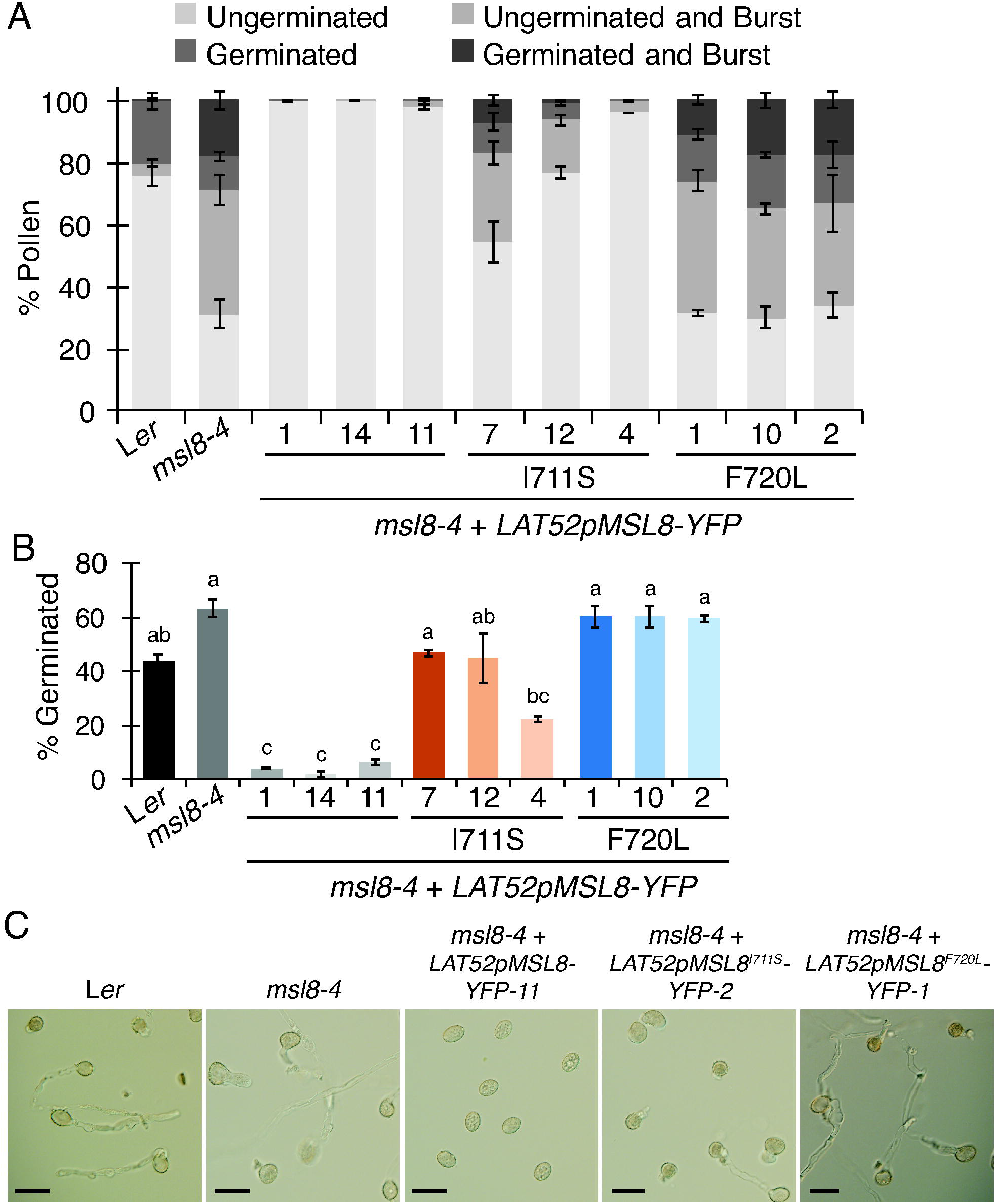
MSL8^I711S^-YFP but not MSL8^F720L^-YFP partially suppresses *msl8-4* pollen bursting and germination when overexpressed in the *msl8-4* background. (**A**) Pollen from the indicated lines incubated in germination media for 6 hours and scored for combined germination and bursting categories as in Fig. 4. Average of 3 to 6 experiments with *N* = 107 to 512 pollen per experiment. See Supplementary Table 2 for statistical differences between groups. (**B**) Germination rate of pollen from the indicated lines after incubation in germination media for 16 hours. Different letter groups indicate significant (p < 0.05) differences between groups as determined by Tukey’s post-hoc test following one-way ANOVA. (**C**) Brightfield images of pollen from the indicated lines after incubation for 16 hours in liquid germination media. Scale bars are 50 μm. (**A**, **B**) Error bars are mean ± SE.

Because germination is suppressed so strongly by *MSL8* overexpression, we also incubated pollen in germination media for an extended period of time (overnight, or 16 hours), in order to maximize the number of germination events (Fig. 7B-C, Supplementary Fig. S2B). Under these conditions, 63% of *msl8-4* pollen germinated on average, compared to 44% of L*er* pollen. The overexpression of *MSL8-YFP* in the *msl8-4* background strongly suppressed pollen germination, while the overexpression of *MSL8^F720L^-YFP* did not. The germination rate of *msl8-4* pollen was partially suppressed by expressing *LAT52pMSL8^I711S^-YFP*, with the strongest-expressing line, I711S-4, germinating 22% of the time and I711S-7 and I711S-12 germinating at rates comparable to L*er*. This indicates that overexpression of MSL8^F720L^ does not rescue the elevated germination and bursting rate of *msl8-4* pollen, and overexpression of MSL8^I711S^ produces a partial reduction in both bursting and germination rates.

We previously observed that overexpressing *MSL8* reduces pollen fertility, likely through the suppression of germination (Hamilton et al., 2015a). This fertility defect can be observed as a reduction in the transmission of the transgene from the hemizygous T1 generation to the segregating T2 generation, as determined by resistance to the herbicide Basta, which is conferred by the transgene. As expected, three lines expressing *LAT52pMSL8-YFP* in the *msl8-4* background exhibited a significant reduction in transgene transmission, reducing Basta resistance from the expected 75% to around 65%. However, none of the lines overexpressing *MSL8^I711S^-YFP* or *MSL8^F720L^-YFP* in the *msl8-4* background showed a significant deviation from the expected rate of resistance (Table 2). This indicates that neither MSL8^I711S^ nor MSL8^F720L^ have an effect on pollen fertility and suggests that the partial suppression of *in vitro* germination conferred by overexpressing *MSL8^I711S^-YFP* in the *msl8-4* background is insufficient to reduce pollen fertility *in vivo*. Thus, while neither MSL8^F720L^ nor MSL8^I711S^ provide clear function in pollen when expressed at endogenous levels, MSL8^I711S^ can provide some function when expressed at high levels.

**Table 2.** Transmission of *LAT52pMSL8-YFP* variant transgenes in the *msl8-4* background.

| *msl8-4* + *LAT52pMSL8-YFP* | T2 progeny resistant to Basta |  |  |  |  |
| --- | --- | --- | --- | --- | --- |
| | Line | % Expected | % Observed<br>(number observed/total) | $\chi^2$ | p |
| *MSL8* | 1 | 75 | 67.2 (86/128) | 5.18 | 0.023 |
|  | 14 | 75 | 66.1 (74/112) | 5.79 | 0.016 |
|  | 11 | 75 | 64.0 (80/125) | 9.52 | 0.002 |
| *MSL8<sup>F720L</sup>* | 1 | 75 | 78.3 (47/60) | 0.23 | 0.64 |
|  | 10 | 75 | 74.4 (29/39) | 0.05 | 0.83 |
|  | 2 | 75 | 75.9 (41/54) | 0.01 | 0.94 |
| *MSL8<sup>I711S</sup>* | 7 | 75 | 74.0 (91/12) | 0.23 | 0.64 |
|  | 12 | 75 | 76.7 (89/116) | 0.07 | 0.79 |
|  | 4 | 75 | 74.1 (86/116) | 0.18 | 0.67 |

### Overexpressing MSL8^F720L^ in wild-type pollen increases bursting, while overexpressing MSL8 and MSL8^I711S^ suppresses pollen germination

We also investigated the function of MSL8^I711S^ and MSL8^F720L^ when overexpressed in the wild type background, rather than in the *msl8-4* background as above. Transgenic *LAT52pMSL8-YFP, LAT52pMSL8^F720L^-YFP*, or *LAT52pMSL8^I711S^-YFP* lines were selected wherein *MSL8* or *MSL8* variants were expressed at levels similar to or higher than the endogenous *MSL8* gene (Fig. 8A). Quantitative RT-PCR revealed that transcript levels of endogenous *MSL8* were not significantly different from L*er* in the transgenic lines (Fig. 8A, dark gray bars), while total *MSL8* transcripts were increased over endogenous levels between 1.3 and 4.3-fold (Fig. 8A, light gray bars).

**Figure 8.**
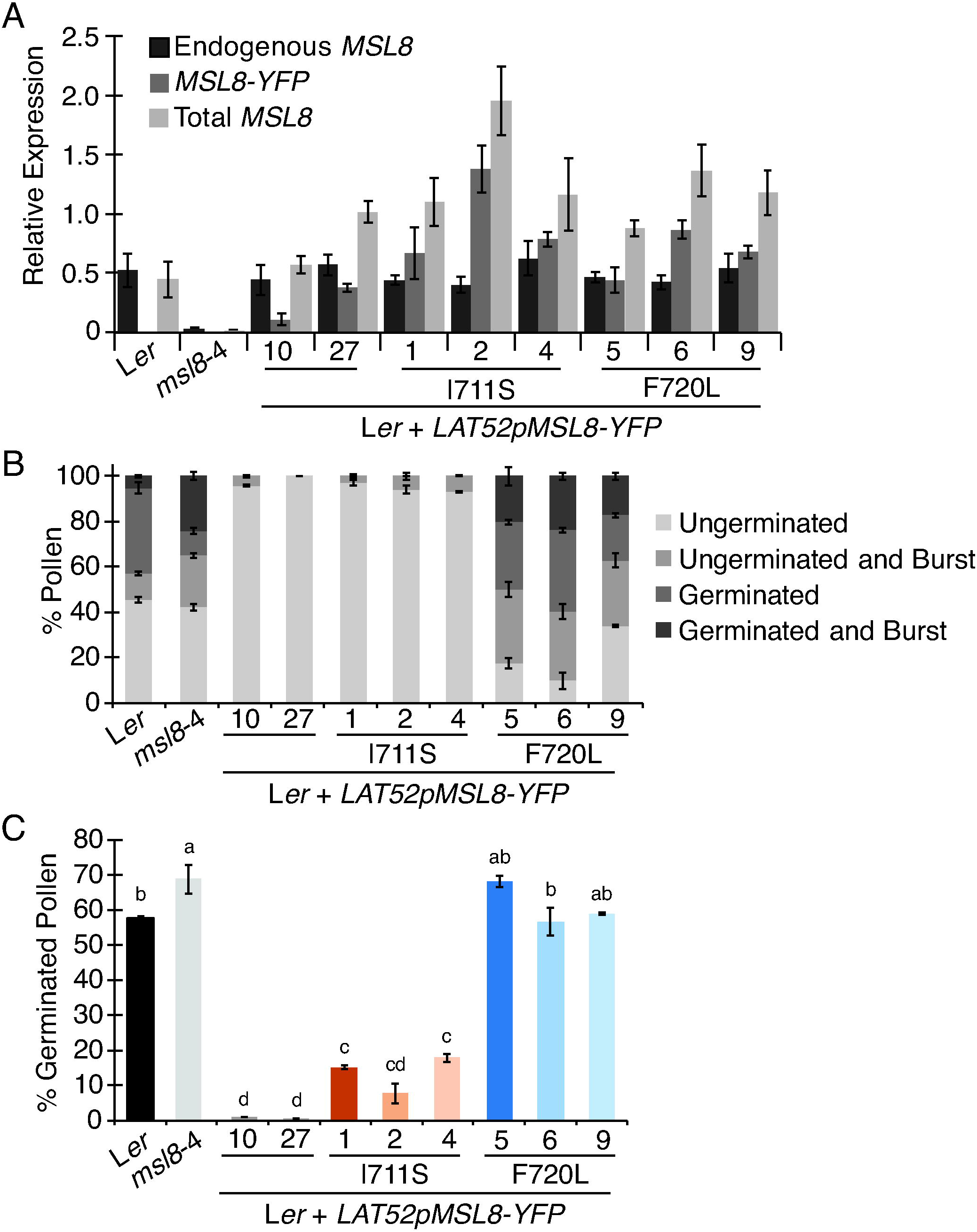
Effect of overexpressing MSL8-YFP variants from the *LAT52* promoter in the wild type background on germination phenotypes. (**A**) Quantitative reverse-transcription polymerase chain reaction of endogenous *MSL8*, *MSL8-YFP*, and total *MSL8* transcripts relative to *ACTIN* in L*er*, *msl8-4*, and *LAT52pMSL8-YFP* transgenic lines. (**B**) Pollen from the indicated lines incubated in germination media for 6 hours and scored for combined germination and bursting categories as in Fig. 4. Averages from 3 experiments with *N* = 82 to 250 pollen per experiment are presented. See Supplementary Table 3 for statistical differences between groups. (**C**) Germination rate of pollen from the indicated lines after incubation in germination media for 16 hours. Averages of 3 experiments with *N* = 99 to 257 pollen per experiment are presented. Different letter groups indicate significant (p < 0.05) differences between groups as determined by Tukey’s post-hoc test following one-way ANOVA. (**A**-**C**) Error bars are mean ± SE.

Pollen from lines expressing *LAT52pMSL8-YFP* and *LAT52pMSL8^I711S^-YFP* did not germinate at all after incubation in germination media for 6 hours, and the bursting rate of ungerminated pollen grains was reduced to less than 7% (Fig. 8B). The pattern of pollen germination and bursting was indistinguishable between line WT-10 and the three lines overexpressing *MSL8^I711S^-YFP* as determined by a chi-squared test (Fig. 8B, Supplementary Table S3). However, L*er* pollen expressing *LAT52pMSL8^F720L^-YFP* burst 46% to 54% of the time, comparable to the bursting rate of *msl8-4*. Thus, overexpressing *MSL8^F720L^* in the wild-type background produced a dominant negative effect, phenocopying the *msl8-4* mutant bursting rate. This elevated bursting rate is likely caused by MSL8^F720L^ disrupting the native pool of MSL8 through the formation of heteromeric channels (see discussion).

When incubated in germination media overnight, pollen overexpressing MSL8^F720L^ in the L*er* background germinated at rates between that of L*er* and *msl8-4*, indicating that it had little, if any, effect (Fig. 8C). However, overexpressing MSL8^I711S^ in the wild type background resulted in a partial suppression of germination rates, to between 9% and 18%. While line I711S-2 had a germination rate that was indistinguishable from the pollen overexpressing wild type MSL8, the other two independent lines produced pollen germination rates that were both significantly lower than that of L*er* and significantly higher than L*er* + *LAT52pMSL8-YFP* lines, representing an intermediate phenotype.

We also quantified the effect of *MSL8* overexpression on male fertility in these lines via the segregation of resistance to Basta. Overexpressing *LAT52pMSL8-YFP* in the L*er* background reduced Basta resistance in the segregating T2 generation from 75% to less than 65% (Table 3). One line expressing *LAT52pMSL8^I711S^-YFP* also exhibited a significant reduction in Basta resistance, to 59%, but all other MSL8 variant lines did not. This indicates that MSL8^F720L^ overexpression does not have an effect on pollen fertility, and suggests that the partial suppression of *in vitro* pollen germination produced by overexpressing *MSL8^I711S^*has only a modest effect *in vivo*.

**Table 3.** **Transmission of *LAT52pMSL8-YFP* variant transgenes in the wild-type background**.

| Ler + *LAT52pMSL8-YFP* | Line | T2 progeny resistant to Basta |  |  |  |
| --- | --- | --- | --- | --- | --- |
| | | % Expected | % Observed<br>(number observed/total) | $\chi^2$ | p |
| *MSL8* | 10 | 75 | 60.0 (30/50) | 6.00 | 0.014 |
|  | 27 | 75 | 63.9 (53/83) | 5.50 | 0.019 |
| *MSL8<sup>F720L</sup>* | 5 | 75 | 78.9 (30/38) | 0.32 | 0.57 |
|  | 6 | 75 | 67.3 (66/98) | 3.06 | 0.080 |
|  | 9 | 75 | 76.9 (40/52) | 0.10 | 0.75 |
| *MSL8<sup>I711S</sup>* | 1 | 75 | 58.6 (34/58) | 8.30 | 0.0040 |
|  | 2 | 75 | 65.6 (40/61) | 2.89 | 0.089 |
|  | 4 | 75 | 72.3 (34/47) | 0.18 | 0.67 |

## Discussion

### Tension-sensitive ion transport activity of MSL8 is critical for its function in pollen

A role for MS ion channels in the response of pollen to mechanical and osmotic challenges was proposed more than 20 years ago (Feijo et al., 1995). Such a role has been solidly established for the canonical MS ion channel from *E. coli*, MscS (Edwards et al., 2005; Levin and Blount, 2004; Levina et al., 1999; Miller, 2003; Sukharev et al., 1994), and an initial analysis suggested that of a MscS homolog from Arabidopsis, MSL8, might as well (Hamilton et al., 2015a). However, it remained possible that MSL8 could play an indirect role in pollen, perhaps signaling independently of ion flux like close homolog MSL10 (Veley et al., 2014). Here we used site-directed mutagenesis, electrophysiology, and physiological assays to determine if the critical function of the MS ion channel MSL8 during pollen hydration and germination is to mediate ion flux.

### Mutating conserved hydrophobic residues in the pore-lining domain of MSL8 resulted in diminished or disrupted channel function

We identified two point mutations that, when introduced into the presumptive pore-lining helix, altered MSL8 channel behavior without appreciably altering expression, stability, or subcellular localization (Figs. 1, 2, Table 1). When expressed in *Xenopus* oocytes, MSL8^I711S^ produced MS currents with a unitary conductance identical to that produced by MSL8 (at pipette potentials from 0-80 mV, 60.7 and. 60.0 pS respectively). However, approximately twice the suction (-29.5 ± 11.7 mmHg for MSL8^I711S^ compared to -18.0 ± 13.1 mmHg for MSL8) was required to open the first MSL8^I711S^ channel compared to the wild type.Thus, MSL8^I711S^ appears relatively insensitive to membrane tension (requiring greater forces to open). When expressed in oocytes, MSL8^F720L^ did produce MS currents that were never observed in water-injected oocytes. However, these MS currents did not form the step-wise increase in current characteristic of individual channel gating events, instead appearing as a flickering increase in current that disappeared immediately after suction was released. Producing these MS currents required higher negative pressures than was required for either MSL8 or MSL8^I711S^. From this we conclude that MSL8^F720L^ is still able to conduct ions through its pore, but is unable to form a stable open state or to transition normally between non-conducting and conducting configurations.

### Disrupting normal channel function prevents MSL8 from complementing known msl8-4 null loss-of-function phenotypes

Our finding that the severely disrupted MSL8^F720L^ channel was unable to rescue *msl8-4* loss-of-function phenotypes either at native levels of expression (Figs. 2-4) or when overexpressed (Figs. 5-7) demonstrated that a functional channel is necessary for MSL8 to protect pollen from the osmotic stress it experiences during hydration and germination. This supports a model where MSL8 functions akin to MscS, acting as an osmotic pressure release valve (Booth and Blount, 2012), albeit during developmentally programmed osmotic challenges rather than in response to environmental swings in osmolarity. During pollen germination and tube growth, which are powered by internal turgor pressure (Benkert et al., 1997; Zerzour et al., 2009), pollen must have multiple mechanisms to ensure that internal pressure and growth rates do not overtake the delivery of cell wall materials to the growing tip or the strength of the cell wall itself (Hill et al., 2012; Kroeger et al., 2011; Zerzour et al., 2009); MSL8 appears to be one of those mechanisms.

### Increasing the tension threshold of the MSL8 channel reduces its physiological activity in pollen

The Ile711Ser mutation increased the force required to gate MSL8, but left wild-type conductance intact– and reduced the ability of MSL8^I711S^ to complement mutant phenotypes. Only when overexpressed could MSL8^I711S^ protect pollen from the hypoosmotic shock of rehydration and maintain the integrity of the pollen tube. Likewise, MSL8^I711S^ only suppressed pollen germination when overexpressed, and not as strongly as the wild type. (Figs. 6-8). These results suggest that wild-type tension sensitivity is crucial for the proper function of MSL8 in pollen. Protection against the hypoosmotic stress of rehydration, maintenance of pollen tube integrity against bursting, and the negative regulation of pollen germination all appear to require that MSL8 gate at tensions below those that open MSL8^I711S^ under native levels of expression.

Although the evidence described above is consistent with MSL8 functioning like an osmotic safety valve, we note that we cannot exclude the possibility that the ion flow across MSL8 could act as a biochemical signal that indirectly achieves the same function.

One example of such biochemical signaling in pollen is the tip-focused Ca^2+^ gradient that regulates pollen tube elongation. High concentrations of Ca^2+^ at the growing pollen tube tip are essential for pollen tube growth (Miller et al., 1992) and require the cyclic nucleotide-gated channel CNGC18 (Gao et al., 2016). Ca^2+^ acts as a crucial secondary messenger in several pathways in order to properly regulate pollen tube growth and guidance to the female gametophyte (Guan et al., 2013). It remains formally possible, though unlikely, that the Ile711Ser or Phe720Leu mutations affect both the characteristics of the channel and the function of other domains important for signaling independent of ion transport. Lesions in both the soluble N-terminus (Veley et al., 2014) and the soluble C-terminus (Zou et al., 2015) affect the cell death signaling function of MSL10, but lesions to the predicted pore-lining domain have not yet been tested. Future work will be required to understand how MSL8 functions alongside established ion channels and the tightly regulated ion fluxes that are essential for pollen germination and tube growth (Michard et al., 2009).

### MSL8^F720L^ has reduced channel activity and can act as a dominant negative allele

The substitution of Leu for Phe at positon 720 in MSL8 both increased the threshold required to produce MS currents and prevented the appearance of the step-wise increases in current that are indicative of a discrete transition from closed to open configurations, suggesting instability in the channel (Fig. 1B, bottom left panel, Supplementary Fig. S1, Table 1). The closed form of *Ec*MscS is proposed to involve close packing of small residues from adjoining pore-lining domains (Edwards et al., 2005). The substitution of large hydrophobic residues for Gly or Ala at these positions results in less stable open state configurations with higher tension thresholds, that at least superficially resemble MSL8^F720L^ (Edwards et al., 2005; Rasmussen et al., 2015; Wu et al., 2011). Alternating chains of small hydrophobic amino acids are not observed in the predicted pore-lining domain of MSL8 (Fig. 1B). Rather, a repeating pattern of large hydrophobic/polar residues in the predicted pore-lining domain is conserved among the seven Arabidopsis MscS homologs predicted to localize to the plasma membrane. These residues include three phenylalanines at positions 710, 720 and 727 (marked with arrowheads, Supplementary Fig. S3). The observed channel characteristics of MSL8^F720L^ suggest that the pairing of similarly sized residues in the pore-lining domains may be an important factor in maintaining the stability and normal function of both MscS and MSLs. Testing this idea will require additional study, in particular detailed structural information for the plasma membrane-localized MSLs.

Expressing *MSL8^F720L^* in a wild type background, and therefore in the presence of a native pool of MSL8, phenocopies the mutant with respect to germination rate and pollen tube bursting (Fig. 8B-C). Because MSLs are likely to form homomeric channels (Haswell et al., 2008; Peyronnet et al., 2008), this dominant negative effect is likely caused by the formation of heteromeric channel complexes. According to this model, the instability caused by the Phe720Leu mutation is dominantly imparted to MSL8-MSL8^F720L^ heteromeric channels, reducing the number of functional MSL8 channels available, and thereby producing pollen with characteristics similar to those from the loss-of-function *msl8-4* background. A similar effect was described for the *Caenorhabditis elegans* TRP-4 candidate MS ion channel (Kang et al., 2010). A TRP-4 mutant that did not form a functional channel on its own dominantly ablated channel function when co-expressed with wild type TRP-4 in touch-sensitive neurons. How many MSL8^F720L^ monomers are sufficient to disrupt the function of an otherwise wild type MSL8 channel is not yet known.

### MSL8^I711S^ can complement the msl8-4 mutant only when overexpressed, consistent with a cooperative gating mechanism

MSL8^I711S^ exhibited both an elevated tension threshold and reduced function *in planta*. MSL8 Ile711 aligns with Ala110 of *Ec*MscS (Fig. 1B), and a mutation in the neighboring residue, Leu111, produces a similar disruption in function. In *Ec*MscS, Leu111Ser is associated with a doubled tension threshold, and MscSL^111S^ is unable to protect *E. coli* against osmotic shock (Belyy et al., 2010). Leu111 is part of the proposed “tension-transmitting clutch” that allows *Ec*MscS to respond to increases in membrane tension through hydrophobic associations with the other TM helices. The substitution of a hydrophilic residue for a hydrophobic one may weaken these interactions. Our data presented here show that Ile711 may play a similar force-transmitting role in MSL8. Once open, MSL8^I711S^ appears to produce a normal pore, as its single-channel conductance was wild type.

There are several possible explanations for the ability of MSL8^I711S^ to partially rescue loss-of-function phenotypes at high levels of expression. First, the gating of a MS ion channel is a stochastic process centered on the average tension threshold (Hille, 1992). Increasing the population of MSL8^I711S^ channels in pollen would increase the number of channels available in the population to open at lower tensions, potentially to levels sufficient for protection against osmotic stress. We do not favor this model, primarily because we do not see a linear increase in complementation associated with expression level. Instead, when expressed at endogenous levels MSL8^I711S^ is completely unable to rescue mutant phenotypes.

Alternatively, MSL8 could participate in cooperative gating. Cooperative gating is the lowering of the average tension threshold as the number of channels embedded in the membrane increases. Biophysical modeling experiments with *Ec*MscL have shown that local deformation of the lipid bilayer at the channel periphery could explain the energetics of cooperative gating (Haselwandter and Phillips, 2013; Ursell et al., 2007). If MSL8^I711S^ is expressed at high enough levels, cooperative gating might reduce the tension threshold enough to allow it to protect pollen during hydration.

Cooperative gating could also explain the observation that overexpressing wild-type MSL8 in the L*er* background suppresses pollen germination (Fig. 8B, (Hamilton et al., 2015a)). It is thought that high turgor pressure is required for breaking through the cell wall during germination (Feijo et al., 1995). If overexpression results in MSL8 channel activity at lower membrane tensions by promoting cooperative gating, then the critical turgor for germination might never be reached. Pollen would then be stuck in a futile cycle of building pressure, relieving pressure, and building it again. In support of this idea, we observed that pollen overexpressing MSL8 developed the larger and more structured vacuole morphology resembling normally germinating pollen (Supplementary Fig. S4; (Hicks et al., 2004; Wudick et al., 2014)), though it rarely went on to germinate. Future work will determine if MSL8 and MSL8^I711S^ exhibit cooperative gating in *Xenopus* oocytes.

### MSL8 fulfills all criteria for assignment as a mechanotransducer

The four criteria necessary for establishing a protein as the transducer of a physiological mechanical response are: (1) it is expressed in the correct cell and subcellular location to respond to mechanical stimulation; (2) it is required for the mechanosensory response, not the normal development of the cell; (3) it forms a MS channel in a heterologous system; and (4) structural changes that affect the protein’s response in a heterologous system affect its function *in vivo* (Arnadottir and Chalfie, 2010). MscL and MscS in bacteria fulfill these criteria (Edwards et al., 2005; Levin and Blount, 2004; Levina et al., 1999; Miller, 2003; Sukharev et al., 1994), and NOMPC of *Drosophila* was recently shown to be a mechanotransducer of touch response in touch-sensitive neurons (Gong et al., 2013; Yan et al., 2013).

MSL8 now fulfills all four criteria: (1) it is expressed in tricellular and mature pollen and pollen tubes, and localizes to the plasma membrane; (2) it is not required for the normal development of pollen, but is required for protection against osmotic stress; (3) it forms a MS channel in the heterologous *Xenopus* oocyte system; and (4) altering its structure changes its electrophysiological characteristics and its physiological function, without affecting expression or subcellular localization.

Pollen faces additional, relatively unstudied mechanical challenges during its development, and mechanically sensitive proteins are likely to be involved. Desiccation is critical for the success of pollen from most species (Franchi et al., 2011), and presents unique osmotic and mechanical challenges (Firon et al., 2012; Hoekstra et al., 1997; 2001). Furthermore, as it invades the sporophytic tissue to reach the female gametophyte, the pollen tube must sense, produce, and regulate the forces required to grow in between other cells (Sanati Nezhad et al., 2013). Finally, the regulated process of bursting that must occur to release the sperm cells from within the pollen tube represents a fascinating case where mechanisms that previously maintained the structural integrity of the cell must be overcome in order to complete fertilization (Amien et al., 2010; Dresselhaus et al., 2016; Woriedh et al., 2013). Using electrophysiology, mutagenesis and physiological assays, we discovered that mechanosensitive ion channels are one mechanism pollen relies on to respond to the mechanical effects of osmotic changes.

This powerful combination of techniques, as well as the development of new tools to probe pollen in the dry state, may uncover other mechanical and ionic regulation strategies in pollen and other plant cells.

## Materials and Methods

### Plant material and growth conditions

Plants were grown on soil under 24 hours of light at 21°C. *msl8-4* (DsLoxN101568 and DsLoxN101751, in the L*er* background) was obtained from the Arabidopsis Biological Resource Center. MSL8^I711S^ and MSL8^F720L^ constructs were produced through site-directed mutagenesis from existing vectors (described in (Hamilton et al., 2015)) and transformed into *msl8-4* or L*er* by floral dip.

### Multiple alignments

Alignments were performed in MEGA7 using a Clustal alignment with a pairwise alignment gap opening penalty of 10 and gap extension penalty of 0.1 and a multiple alignment gap opening penalty of 10 and gap extension penalty of 0.2. Sequences for alignment were based on previous phylogenetic analysis of *Ec*MscS and Arabidopsis MSLs (Haswell, 2007) predicting the pore-lining domains of MSLs based on the known pore-lining domain of *Ec*MscS.

### Electrophysiology

Single channel patch-clamp electrophysiology was performed as described in (Maksaev and Haswell, 2015). Capped RNA (cRNA) of *MSL8* was transcribed *in vitro* using the mMESSAGE mMACHINE SP6 kit (Ambion) and stored at 1µg/µl at -80 °C. Defolliculated oocytes were purchased from Xenopus1 (Dexter, MI) and injected with 50 nl of cRNA or water and patched in symmetric 60 mM MgCl2 after incubating for 2-10 days in ND96 buffer + gentamycin.

### Reverse-transcriptase-polymerase chain reaction

RNA was isolated from floral tissue (stage 13/14 flowers) using the Qiagen RNeasy Mini RNA extraction kit (Qiagen). cDNA was generated from 2 µg RNA using an oligo(dT)20 primer and the M-MLV Reverse Transcriptase kit (Promega). *ACTIN,* total *MSL8,* endogenous *MSL8* and *MSL8-YFP* transcripts were amplified with the primers listed in Supplementary Table S4 using SYBR Green PCR Master Mix (Applied Biosciences) and 0.25 µL cDNA at a final volume of 25 µL. Quantitative RT-PCR was performed using the StepOnePlus real-time PCR system (Applied Biosystems). Total *MSL8* was amplified using primers that do not distinguish between the endogenous locus and the transgenes. Endogenous *MSL8* was amplified using a reverse primer in the 3’ UTR, which is not present in the transgenes.

### In vitro pollen hydration

Pollen from mature (stage 13-14) flowers was hydrated in 25-30 μl drops of water or the indicated percentage of PEG (average molecular weight 3350 g/mol, Sigma-Aldrich) at a final concentration of 1 μg/ml fluorescein diacetate (FDA, Sigma-Aldrich) and 0.5 μg/ml propidium iodide (PI, Sigma-Aldrich) on double-ring cytology slides. Slides were inverted and incubated in a humid chamber at room temperature for the indicated amount of time. To image, cover slips were added and FDA signal was collected in the GFP epifluorescence channel while PI signal was collected in the dsRed epifluorescence channel. FDA stains live pollen while PI enters dead pollen.

### In vitro pollen germination

Pollen germination was performed according to (Daher, Chebli, Geitmann 2009 Plant Cell Rep). Pollen was pre-hydrated by removing flowers from plants and incubating for 45 minutes at 30°C in a humid chamber constructed from a large petri dish containing smaller petri dishes placed on top of moistened filter paper. Pollen was incubated in 30 μl of pollen germination media (2 mM CaCl2, 2 mM Ca(NO3)2, 0.49 mM H3BO3, 1 mM MgSO4, 1 mM KCl, 18% w/v sucrose, pH 7) at 30°C for 6 or 16 hours in a humid chamber as during pollen hydration. Pollen was counted as germinated if it had produced a pollen tube longer than the pollen grain. Pollen was counted as burst if expelled cytoplasm was visible outside the pollen grain or pollen tube.

### Microscopy

Confocal images of GFP or YFP signal in pollen were acquired on an Olympus BX-61 microscope using FV10-ASW Olympus software and the GFP (488 nm excitation, 505-605 nm bandpass filter) or YFP (515 nm excitation, 535-565 nm bandpass filter) channels. Brightfield and epifluorescent images for pollen germination and pollen viability assays were collected on the same microscope using an Olympus DP71 digital camera, DP Controller software, and filter sets for GFP (470/40 nm excitation, 525/50 nm emission) or dsRED (545/30 nm excitation, 620/60 nm emission).

### Calculation of transmission ratios

The transmission frequencies of the *MSL8* transgenes were determined by selecting seedlings from the T2 generation with Basta on soil and counting the number of sensitive and resistant progeny. The Basta resistance gene *bar* is included in the transgene.

### Statistical analyses

One-way or two-way ANOVAs were performed as indicated in figure legends. Tukey’s HSD post-hoc test was used to determine statistical significance for balanced data sets. Scheffe’s post-hoc test was used to determine statistical significance for unbalanced data sets. Chi-squared tests with Bonferroni correction were performed for analysis of transgene transmission and for comparison between groups of the pattern of germinated and burst pollen during pollen germination assays.

## Acknowledgements

We thank G. Maksaev for advice and assistance. We acknowledge the Washington University plant growth facility for their assistance.

## Abbreviations

cRNA: capped RNA
MS: mechanosensitive
MscL: Mechanosensitive channel of Large conductance
MscS: Mechanosensitive channel of Small conductance
MSL8: MscS-Like 8
MSL10: MscS-Like 10
PEG: polyethylene glycol
TM: transmembrane

